# Discordant bioinformatic predictions of antimicrobial resistance from whole-genome sequencing data of bacterial isolates: An inter-laboratory study

**DOI:** 10.1101/793885

**Authors:** Ronan M. Doyle, Denise M. O’Sullivan, Sean D. Aller, Sebastian Bruchmann, Taane Clark, Andreu Coello Pelegrin, Martin Cormican, Ernest Diez Benavente, Matthew J. Ellington, Elaine McGrath, Yair Motro, Thi Phuong Thuy Nguyen, Jody Phelan, Liam P. Shaw, Richard A. Stabler, Alex van Belkum, Lucy van Dorp, Neil Woodford, Jacob Moran-Gilad, Jim F. Huggett, Kathryn A. Harris

## Abstract

**Background:** Antimicrobial resistance (AMR) poses a threat to public health. Clinical microbiology laboratories typically rely on culturing bacteria for antimicrobial susceptibility testing (AST). As the implementation costs and technical barriers fall, whole-genome sequencing (WGS) has emerged as a ‘one-stop’ test for epidemiological and predictive AST results. Few published comparisons exist for the myriad analytical pipelines used for predicting AMR. To address this, we performed an inter-laboratory study providing sets of participating researchers with identical short-read WGS data sequenced from clinical isolates, allowing us to assess the reproducibility of the bioinformatic prediction of AMR between participants and identify problem cases and factors that lead to discordant results.

**Methods:** We produced ten WGS datasets of varying quality from cultured carbapenem-resistant organisms obtained from clinical samples sequenced on either an Illumina NextSeq or HiSeq instrument. Nine participating teams (‘participants’) were provided these sequence data without any other contextual information. Each participant used their own pipeline to determine the species, the presence of resistance-associated genes, and to predict susceptibility or resistance to amikacin, gentamicin, ciprofloxacin and cefotaxime.

**Results:** Individual participants predicted different numbers of AMR-associated genes and different gene variants from the same clinical samples. The quality of the sequence data, choice of bioinformatic pipeline and interpretation of the results all contributed to discordance between participants. Although much of the inaccurate gene variant annotation did not affect genotypic resistance predictions, we observed low specificity when compared to phenotypic AST results but this improved in samples with higher read depths. Had the results been used to predict AST and guide treatment a different antibiotic would have been recommended for each isolate by at least one participant.

**Conclusions:** We found that participants produced discordant predictions from identical WGS data. These challenges, at the final analytical stage of using WGS to predict AMR, suggest the need for refinements when using this technology in clinical settings. Comprehensive public resistance sequence databases and standardisation in the comparisons between genotype and resistance phenotypes will be fundamental before AST prediction using WGS can be successfully implemented in standard clinical microbiology laboratories.

## Introduction

Antimicrobial resistance (AMR) is a major, global, public health threat with projections of up to 10 million deaths per annum by 2050 [1]. The World Health Organisation’s 2015 Global Action Plan on AMR identified diagnostics as a priority area for combating resistance [2]. Whilst most diagnostic AMR testing is phenotypic antimicrobial susceptibility testing (AST) based on principles dating back to the early 20^th^ century [3]. Molecular testing has facilitated the implementation of PCR assays that target key AMR mutations and genes [4,5]. However there remains an unmet need for truly rapid point-of-care AST [6,7].

Whole-genome sequencing (WGS) is emerging as a routine clinical test that could be used to determine the bacterial species, undertake transmission tracking and identify multiple AMR associated mutations and genes in a single assay [8–13]. Whilst the initial clinical roll-out of WGS has used bacterial isolates, metagenomics and sequencing direct from clinical samples are future possibilities [14–16]. Resolving the challenges of AMR prediction using WGS for bacteria will provide key advances for the application of metagenomics as a clinical test.

Bioinformatics tools and pipelines to predict AMR have generally been developed by individual research groups, many with no clinical expertise, and mostly with the same basic principle of matching the input DNA sequence to entries in a reference database of known AMR-associated gene sequences. The testing of pipelines for AMR prediction is typically either performed in house [17–19] or done *ad hoc* for specific research [20–23]. Often, these tools are not developed with clinical application or portability in mind. Currently there are no higher-order reference materials (synthetic references that contain exact components of interest) that are available to validate these tools. Studies have reported good concordance between genotype and phenotype on datasets they have been applied to [9,21,24], but rarely address the factors underlying situations where different methods may produce discordant results and how this discordance should be resolved.

Gaining laboratory accreditation is an important, often essential step for tests in clinical microbiology, but is less advanced for bioinformatics due to its comparatively recent development. Bioinformatic reproducibility studies have been performed for clinically relevant bacterial sequence typing methods [25,26]. However, while there have been intra-laboratory studies comparing methods of AMR prediction, there have been no comparisons of multiple methods at the inter-laboratory scale. As there is limited evidence of robust, reproducible analyses in bioinformatic prediction of AMR from clinical WGS data, adoption of these methods may be hampered in meeting the necessary accreditation.

This multi-centre study used genomic DNA sequences from clinical carbapenem-resistant organisms (CROs), specifically chosen to be of varying quality and complexity, to identify the contributors to discordant AMR predictions. Participants included a mixture of individuals and teams involved in AMR prediction from research groups, hospital laboratories, public health laboratories and clinical diagnostic companies. The observations made underpin our recommendations for future method developments.

## Methods

### Sample collection and whole genome sequencing

For the purposes of this study, a panel of ten samples (A-1, A-2, B-1, B-2, C-1, C-2, D, E, F and G) were generated from seven clinical isolates (A, B, C, D, E, F and G). The bacteria were isolated between 2014 and 2017 from stool specimens from patients attending Great Ormond Street Hospital (GOSH) UK or University Hospital Galway (UHG), Ireland. They represented six clinically-relevant bacterial species, including diverse Enterobacterales and also *Acinetobacter baumannii*, and contained six distinct families of carbapenemase genes (Table 1).

**Table 1.** Inter-laboratory study sample characteristics.

| Study ID | Isolate species | Sequencing method | Carbapenemase gene | Median depth of coverage | Comment |
| --- | --- | --- | --- | --- | --- |
| A-1 | *K. pneumoniae* | NEBNext Ultra II + NextSeq 150bp PE | OXA-48-like | 190.2 | Exact duplicate of A-2 |
| A-2 | *K. pneumoniae* | NEBNext Ultra II + NextSeq 150bp PE | OXA-48-like | 190.2 | Exact duplicate of A-1 |
| B-1 | *E. cloacae* complex | NEBNext Ultra II + NextSeq 150bp PE | OXA-48-like | 1.4 | Very low coverage duplicate of B-2 |
| B-2 | *E. cloacae* complex | NEBNext Ultra II + NextSeq 150bp PE | OXA-48-like | 142.9 | High coverage duplicate of B-1 |
| C-1 | *K. oxytoca* | Nextera DNA + HiSeq 100bp PE | OXA-48-like | 37.4 | Same original isolate as C-2 |
| C-2 | *K. oxytoca* | NEBNext Ultra II + NextSeq 150bp PE | OXA-48-like | 156.4 | Same original isolate as C-1 |
| D | *K. pneumoniae* | NEBNext Ultra II + NextSeq 150bp PE | NDM | 83.5 |  |
| E | *E. coli* | Nextera DNA + HiSeq 100bp PE | IMP | 20.6 |  |
| F | *C. freundii* | NEBNext Ultra II + NextSeq 150bp PE | VIM | 32.5 |  |
| G | *A. baumannii* | NEBNext Ultra II + NextSeq 150bp PE | OXA-23-like & OXA-51-like | 22.2 |  |

Phenotypic AST was performed at UHG and GOSH using the EUCAST disk diffusion method (http://www.eucast.org) and meropenem, ertapenem, cefotaxime, amikacin, gentamicin and ciprofloxacin. The isolates were confirmed as carbapenemase producers by PCR at a reference laboratory (Public Health England).

Total genomic DNA was extracted from isolate sweeps on the EZ1 Advanced XL (Qiagen) using DNA Blood 350 µl kits with an additional bead beating step. For eight samples the NEBNext Ultra II DNA Library Prep Kit (New England Biolabs) and NextSeq (Illumina) 150bp paired-end sequencing was used. For two samples Nextera DNA Library Prep Kit (Illumina) and HiSeq 100bp paired-end sequencing was used (Table 1). The FASTQ files were deposited in the European Nucleotide Archive (PRJEB34513).

### Inter-laboratory study plan

Potential inter-laboratory participants were invited in an individual capacity both in person and by email at the meeting “Challenges and new concepts in antibiotics research”, March 2018, at Institut Pasteur, France. Fifteen individuals were also emailed directly to participate in the study. From those invited, nine sets of participants agreed to take part in the study. We will refer to these sets as ‘participants’ throughout. These participants were labelled Lab_1 to Lab_9; “Lab” is used as a catch-all term for an individual or team of participants, who came from a mixture of research groups, hospital laboratories, public health laboratories, and clinical diagnostic companies. All participants agreed to take part in a personal capacity under the condition of anonymity of the results. Each participant was not made aware who the other invited participants were at this stage.

Participants were sent ten paired FASTQ files (labelled AMRIL_1 to AMRIL_10) and were blinded to their contents. The samples included: Two exact duplicates A-1 and A-2 (renamed copies of the same FASTQ files). Two duplicates with different sequence coverage B-1 and B-2 (sequenced from the same isolate, but with median read depths of 1.4X and 142.9X respectively). Two samples sequenced from the same isolate C-1 and C-2 (sequenced in two different laboratories using NextSeq and HiSeq respectively). The remaining four samples D, E, F and G represented diverse bacterial species and carbapenemases).

Participants were asked to report a species identification for each pair of FASTQ files provided as well as the presence of all AMR-associated genes present in that sample. They were asked, using the above data, to make a categorical prediction on whether that sample would be resistant to ciprofloxacin, gentamicin, amikacin and cefotaxime. Lastly, participants were asked to provide a detailed description of the analysis pipeline they used.

Participants returned results via an Excel spreadsheet (Additional file 1). Results were collated for all species identification and resistant or susceptible predictions from each participant. Collated AMR-associated genes had each name manually checked between each participant to identify minor differences in nomenclature used. Individual methods are summarised in Table 2. The full methods submitted by each participant can be found in Additional file 2.

**Table 2.** Summary of bioinformatic tools used for detecting antimicrobial resistance by each participant.

| Method step | Lab_1a<sup>1</sup> | Lab_1b<sup>1</sup> | Lab_2 | Lab_3 | Lab_4 | Lab_5 | Lab_6 | Lab_7 | Lab_8 | Lab_9 | References |
| --- | --- | --- | --- | --- | --- | --- | --- | --- | --- | --- | --- |
| Read assembly | shovill (SPAdes) | shovill (SPAdes) | SPAdes | Unicycler (SPAdes) | No assembly | A5-miseq | Bionumerics | No assembly | Unicycler (SPAdes) | No assembly | [27–29] |
| AMR identifier | RGI | c-SSTAR | ABRicate | RGI & Resfinder | ARIBA | RGI | Bionumerics *E. coli* genotyping plugin (BLAST) | SRST2 | ABRicate | Genefinder | [17,19,30–32] |
| Reference database | CARD | Resfinder & ARG-ANNOT | CARD | CARD & Resfinder | CARD & ARG-ANNOT | CARD | Resfinder | ARG-ANNOT | Resfinder | CARD & Resfinder (manually curated) | [17,31,33] |
| Sequence identity cut-off | 80% | 95% | 75% | 80% (CARD) & 90% (Resfinder) | 90% | 80% | 90% | 90% | 75% | 90% |  |
| Sequence coverage cut-off | 0% | 0% | 0% | 0% (CARD) & 80% (Resfinder) | 20% | 0% | 60% | 90% | 0% | 100% |  |

1. Lab\_1 provided two sets of results with two separate methods and so have been split into Lab\_1a and Lab\_1b.

## Results

### Bacterial species identification

Four of the nine participants identified all species correctly from WGS data (Table 3). This included sample B-1 where we did not expect enough information for a correct call. Species misidentifications of D and B-2 at the genus level by Lab_5 is likely to be human reporting error rather as they correctly identified species in B-1 from a very low read depth. Lab_6 used the same web-based tool for species identification as Lab_5 (Kmerfinder, CGE) but one error was noted where raw sequence reads were inputted instead of assembled contiguous sequences (Table 3).

**Table 3.**
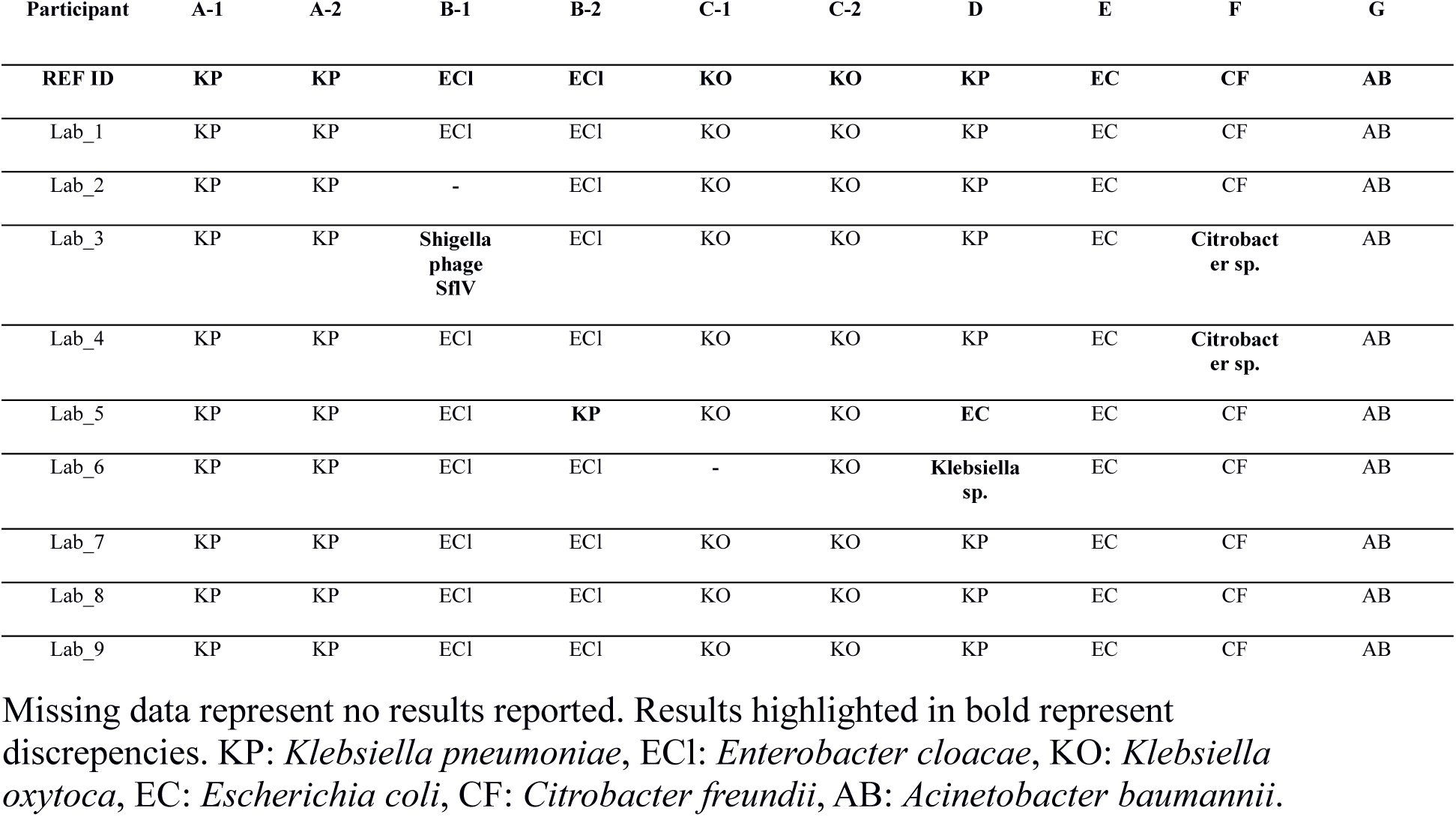
Species identification for each sample by each participant.

### Antimicrobial resistance gene identification

We compared the number of AMR-associated genes reported by each participant in each sample and found disparities in the total reported (Figure 1). Lab_1 used two different methodologies for identifying AMR-associated genes; these are referred to as Lab_1a and Lab_1b. The number of AMR-associated genes reported by each participant was affected by the choice of database used. Lab_1a, Lab_2, Lab_3 and Lab_5 all repeatedly reported the highest number of genes in each sample and all used the Comprehensive Antibiotic Resistance Database (CARD) as their reference database. This is due to CARD including many sequences from loosely AMR-associated efflux pump genes that are not found in the other databases. Lab_4 and Lab_9 also used CARD but in combination with other databases and selectively reported genes. The number of AMR-associated genes reported by each participant was also found to be associated with sequence identity and coverage thresholds used to infer a “hit”. Both Lab_2 and Lab_8 used the lowest identity and coverage thresholds (75% sequence identity and no coverage threshold) and lab_2 consistently reported the highest number of AMR genes in each sample. While Lab_8 reported fewer AMR-associated genes than Lab_2, it did use ResFinder as its reference database rather than CARD, and reported the highest number of genes compared with other participants using the same database.

**Figure 1.**
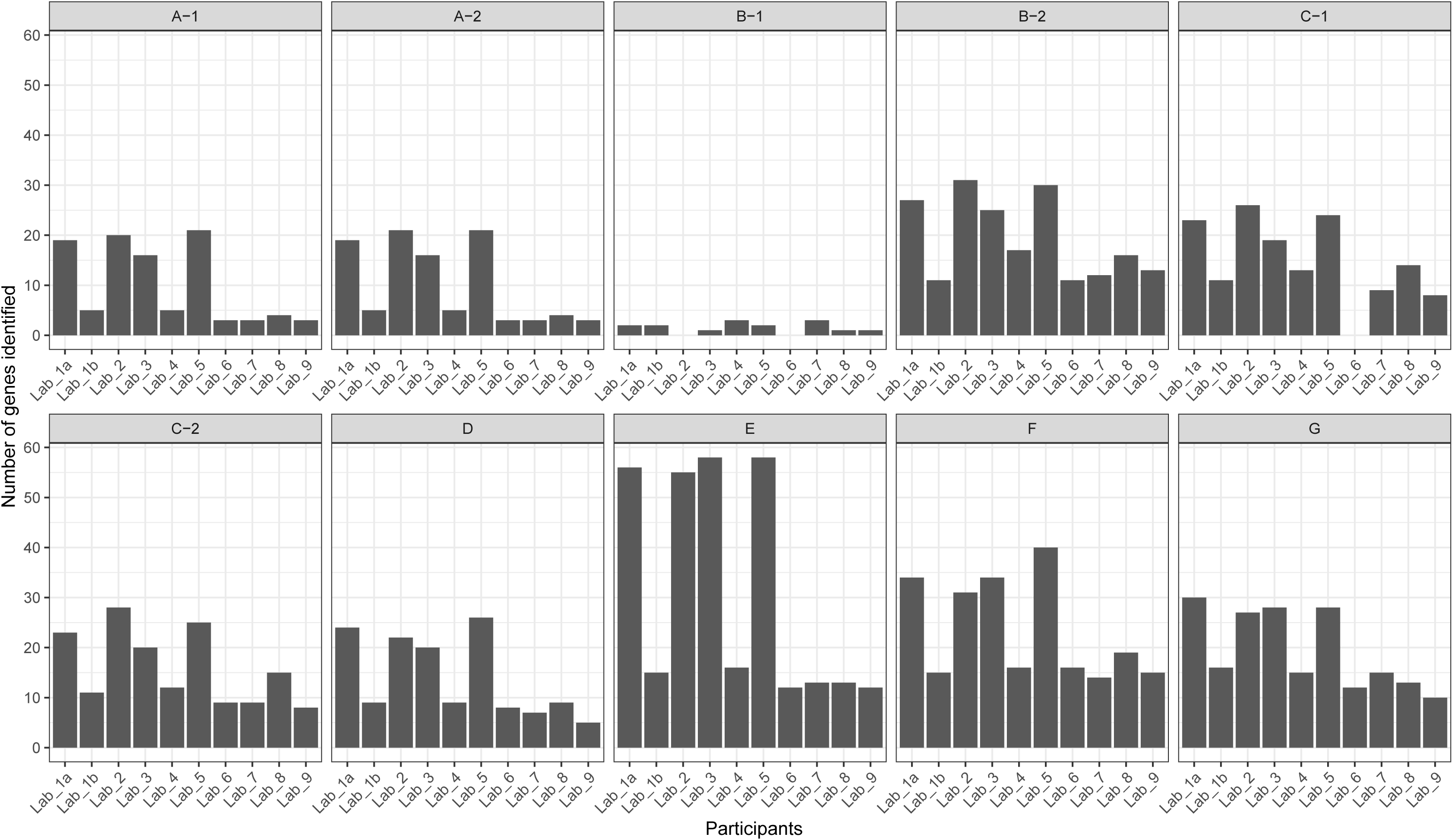
Number of antimicrobial resistance associated genes identified in each sample.

All isolates included in this study were carbapenem resistant. The reporting of carbapenemase genes from whole-genome sequencing from all participants matched the reference PCR result in 91% of cases (91/100) (Table 4). Eight of the ten misidentifications occurred in the low coverage sample B-1 as would be expected. Differences between reported gene variants of *bla*_IMP_ were seen in sample E. Five participants reported *bla*_IMP-1_, whereas the other five reported *bla*IMP-34. This discrepancy exactly matched the reference database used with those reported *bla*_IMP-1_ having used CARD and those who reported *bla*_IMP-34_ either having used ResFinder or ARG-ANNOT. While the sequences for *bla*_IMP-34_ included in each database are identical, the choice of *bla*_IMP-1_ reference sequence included in both databases only share 85% sequence identity. This is due to CARD’s *bla*_IMP-1_ reference sequence being isolated from a *Pseudomonas aeruginosa* integron (NCBI accession: AJ223604) and ARG-ANNOT’s reference sequence from a *Acinetobacter baumannii* integron (NCBI accession: HM036079). While there is variation at the nucleotide level, both encode the same IMP-1 enzyme.

**Table 4.** Carbapenemase genes identified for each sample by each participant and the reference laboratory PCR.

| Participant | A-1 | A-2 | B-1 | B-2 | C-1 | C-2 | D | E | F | G |
| --- | --- | --- | --- | --- | --- | --- | --- | --- | --- | --- |
| REF PCR<sup>1</sup> | OXA-48-like | OXA-48-like | OXA-48-like | OXA-48-like | OXA-48-like | OXA-48-like | NDM | IMP | VIM | OXA-23-like + OXA-51-like |
| Lab_1a<sup>2</sup> | OXA-48 | OXA-48 |  | OXA-48 | OXA-181 | OXA-181 | NDM-1 | IMP-1 | VIM-4 | OXA-23 + OXA-66 |
| Lab_1b<sup>2</sup> | OXA-48 | OXA-48 | OXA-48 | OXA-48 | OXA-181 | OXA-181 | NDM-1 | IMP-34 | VIM-4 | OXA-23 + OXA-66 |
| Lab_2 | OXA-48 | OXA-48 |  | OXA-48 | OXA-181 | OXA-181 | NDM-1 | IMP-1 | VIM-4 | OXA-23 + OXA-66 |
| Lab_3 | OXA-48 | OXA-48 |  | OXA-48 | OXA-181 | OXA-181 | NDM-1 | IMP-1 | VIM-4 | OXA-23 + OXA-66 |
| Lab_4 | OXA-48 | OXA-48 |  | OXA-48 | OXA-181 | OXA-181 | NDM-1 | IMP-34 + IMP-9 | VIM-4 | OXA-23 + OXA-66 |
| Lab_5 | OXA-48 | OXA-48 |  | OXA-48 | OXA-181 | OXA-181 | NDM-1 | IMP-1 | VIM-4 | OXA-23 |
| Lab_6 | OXA-48 | OXA-48 |  | OXA-48 | OXA-181 | OXA-181 | NDM-1 | IMP-34 | VIM-4 | OXA-23 + OXA-66 |
| Lab_7 | OXA-48 | OXA-48 | OXA-48 | OXA-48 | OXA-181 | OXA-181 | NDM-1 | IMP-34 | VIM-4 | OXA-23 + OXA-66 |
| Lab_8 | OXA-48 | OXA-48 |  | OXA-48 | OXA-181 | OXA-181 | NDM-1 | IMP-34 | VIM-4 | OXA-23 + OXA-66 |
| Lab_9 | OXA-48 | OXA-48 | OXA-405 | OXA-48 | OXA-181 | OXA-181 | NDM-1 | IMP-1 | VIM-4 | OXA-23 + OXA-66 |

1. Specific carbapenemase PCR results for each sample.
2. Lab\_1 provided different results using two separate methods and so are included as Lab\_1a and Lab\_1b.
Missing data represent no results reported.

We compared all AMR-associated genes identified by each participant in each sample. As previously noted, the largest discrepancy were the 55 efflux pump gene sequences which were present only in CARD (Figure S1). To understand the other factors influencing discordant reporting we removed these genes that were only present in one database from our comparisons (Figure 2). A pairwise comparison between all participants found that two participants only reported the exact same genes within a sample in 2% (18/900) of cases. Fourteen of these cases occurred when analysing the two identical samples (A-1 and A-2, Figure 2). Although there was little agreement between participants for genes identified in A-1 and A-2, there was complete within-participant concordance across both samples, exhibiting reproducibility within each analysis pipeline. No two participants reported the exact same combination of gene variants in samples B-2, C-1, D, F and G. There were many clear examples where participants assigned different gene variants to the same sequence data where the reference sequences only differed by a few single nucleotides. This can be seen in Figure 2 amongst samples which contained tetracycline resistance genes (*tet(A), tet(B)* and *tet(C)*), some aminoglycoside modifying enzyme gene variants (*aac(3)-IIa* and *aac(3)-IIc*) and *β*-lactamases (*bla*_ACT-14_ and *bla*_ACT-18_). We also observed differences between the same participants analysing samples from the same original isolate. Due to the very low read depth, the genes reported in B-1 bore little resemblance to B-2 across all participant results. However even in the samples from the same isolates with sufficient sequencing depth (C-1 and C-2) we observed differences in the genes identified in four out of nine participants. This suggests that resequencing, and even small increases in read length, can produce variation in results. It is worth noting that all but one of these differences were additional genes identified in C-2, which had a higher read depth than B-2 (156 vs 37 median read depth). The additional genes in C-2 included *ant(3’’)-Ia* (lab_2 and lab_8), *fosA7* (lab_2 and lab_8) and *tet(C)* (lab_3) but the reported reference coverage of *ant(3’’)-Ia* and *fosA7* was low (17% and 75%, respectively) and the sequence similarity between the purported *tet(C)* sequence and the reference was also low (75%). We also found no systematic differences in genes present or absent between those participants that used tools that required assembly of short reads first and those that took unassembled short reads as input (lab_4 and lab_8, ARIBA and SRST2 respectively).

**Figure 2.**
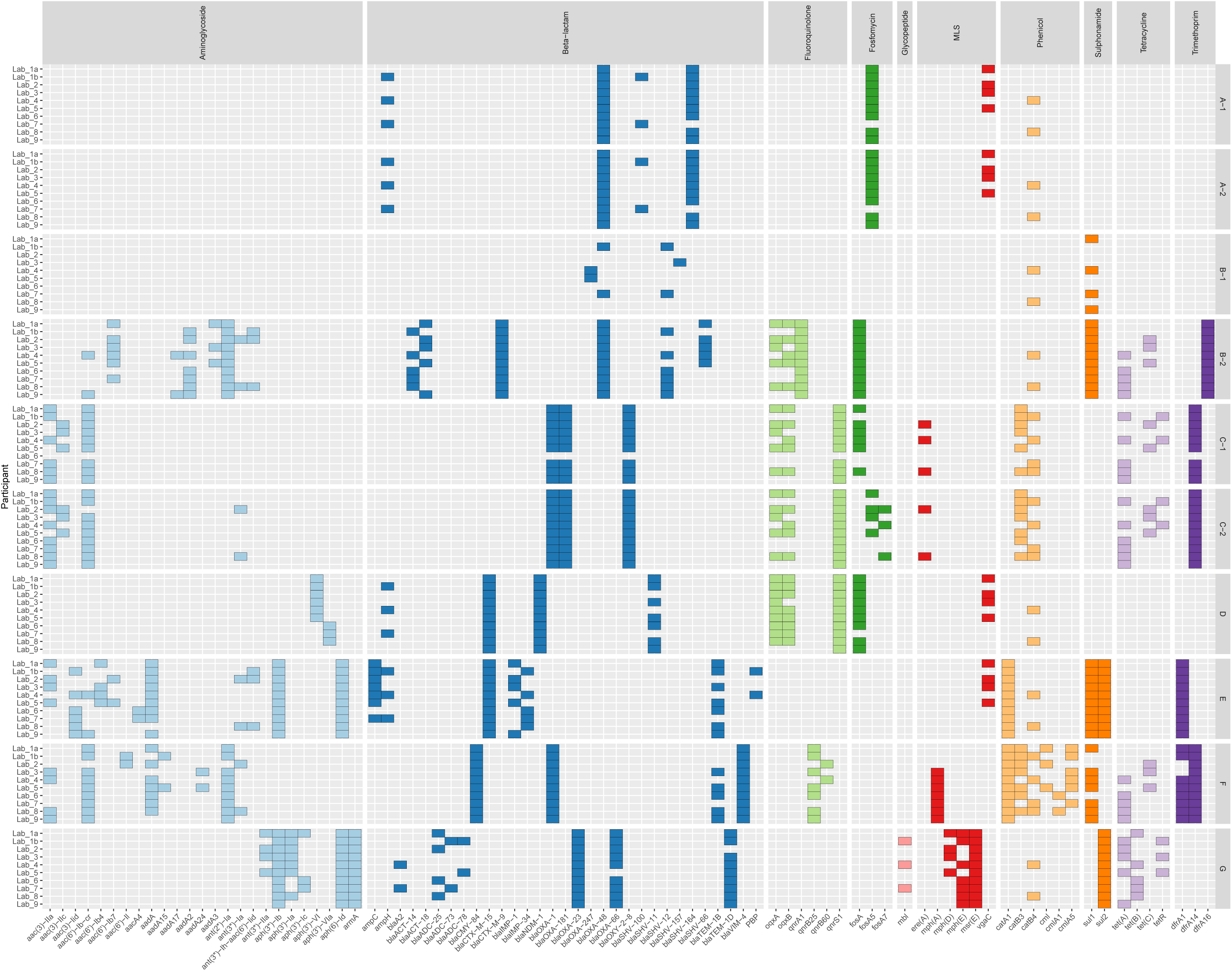
Presence of AMR-associated genes in each sample by each participant. Genes are organised and coloured by the class of antibiotics they are associated with resistance. Genes are only shown here if reported by more than one participant and if they were present in more than one reference database used.

### Phenotypic and genotypic resistance concordance

Given the differences in the AMR-associated genes identified in the samples by each participant, we also compared predictions of antibiotic resistance to phenotypic AST results and each other. Two participants (Lab_2 and Lab_4) did not submit any results for phenotypic resistance prediction and so were not included in the subsequent analysis. A pairwise comparison between genotypic prediction results reported by all participants, on all antibiotics and samples, showed an overall consensus of 79% (864/1092, Figure 3). This varied depending on the antibiotic tested with the highest pairwise reporting consensus of 88% (240/273) between participants for ciprofloxacin and the lowest pairwise reporting consensus of 72% (197/273) for cefotaxime, which could be understandable given the different complexities of the resistance mechanisms involved. When we compared results from each participant with the phenotypic AST results, we found an overall sensitivity of 76% and specificity of 50%. Broken down by antibiotic, the highest consensus between phenotype and genotype was gentamicin (78%, 62/79) and the lowest amikacin (43% 34/79). As expected, there was little agreement between predictions within the low read depth sample (B-1) and most participants predicted a susceptible isolate due to missing data when in fact it was resistant by phenotypic AST. However, when analysing the same isolate at a higher read depth (B-2) there was near perfect concordance between participant reported genotypes and the resistance phenotype, with only two discrepant results reported by Lab_3 (ciprofloxacin) and Lab_7 (amikacin). Lab_3 also reported different results between the two identical samples (A-1 and A-2) where A-1 was reported as resistant and A-2 was reported as sensitive. As there were no differences in the gene content reported in either sample by this participant (Figure 2), this is likely to be due to human reporting error. We also identified a single discrepancy between amikacin resistance predicted by Lab_7 between samples C-1 and C-2 which both were sequenced from the same isolate. C-1 was reported as sensitive but C-2 was reported as resistant and the phenotypic AST result was sensitive, however there was no difference in the reported gene content in both samples by Lab_7 so it is also another likely human reporting error. Excluding the extremely low depth sample, B-1, there were only 2/30 cases where no laboratory correctly predicted the phenotypic AST result. Both of these results were an incorrect resistance prediction for amikacin in C-2 and E but as noted earlier the prediction from Lab_7 for C-2 was likely human error.

**Figure 3.**
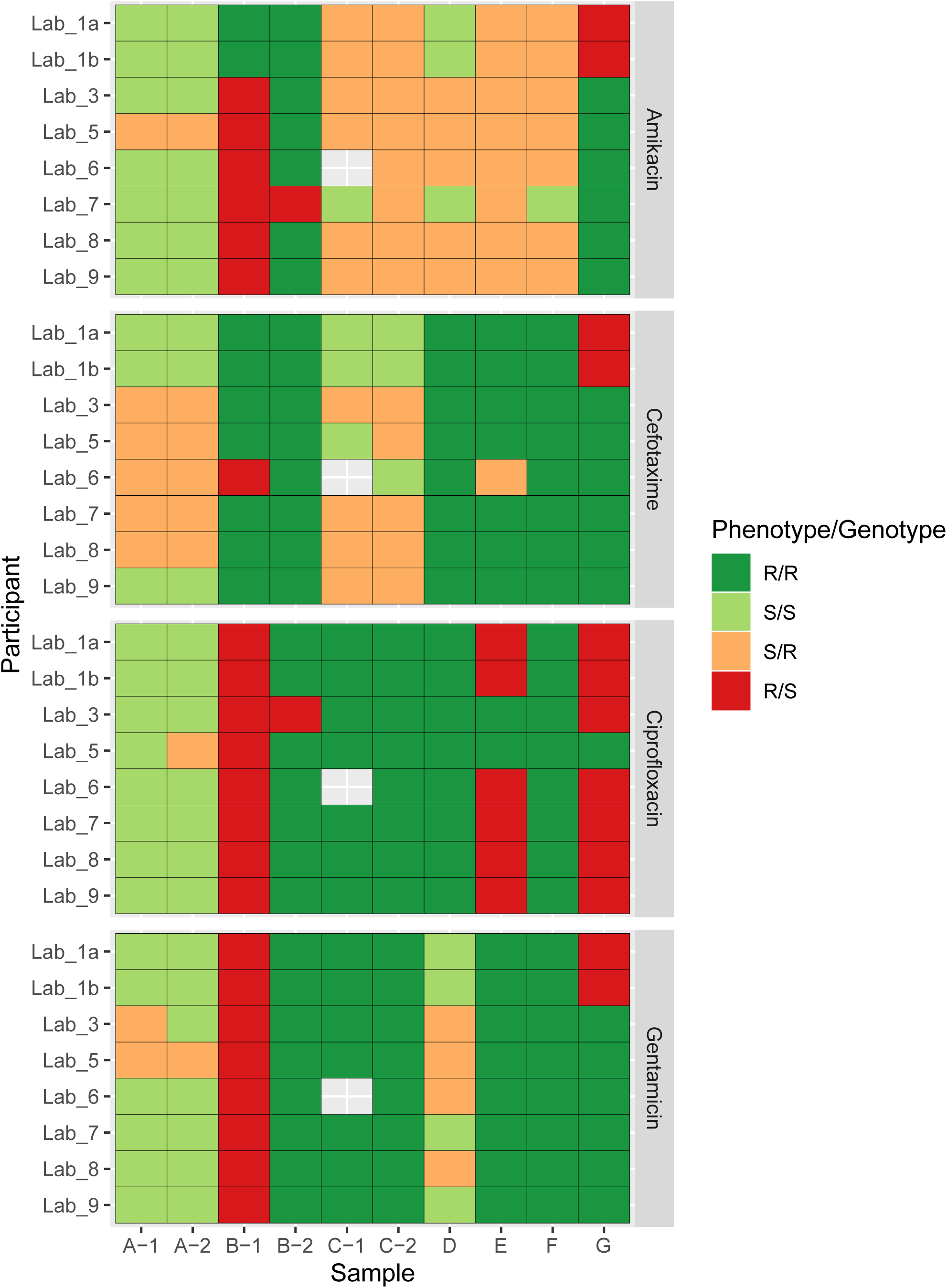
Concordance between phenotypic AST result and the genotypic prediction from WGS data. Results are presented separately for each participant sample and antibiotic. Each tile is coloured based on whether both the resistant phenotype and genotype agreed (R/R). Both phenotype and genotype predicted sensitive (S/S). Major errors where the phenotype was sensitive but the genotype was resistant (S/R) and very major errors where the phenotype was resistant but the genotype was sensitive (R/S). Missing cells represent a result not reported.

## Discussion

In this study we have shown that participants using different bioinformatics pipelines report different AMR-associated gene variants when given identical bacterial isolate WGS datasets and that this led to differences in reporting of predicted resistance phenotypes. We observed good concordance for genotypic resistance predictions between participants but poor concordance with phenotypic AST results. A similar trend has previously been seen in a study of *Staphylococcus aureus* genomes [34]. Concordance in phenotype prediction differed for different antibiotic classes. Good concordance was seen comparing WGS with AST results for gentamicin, but for amikacin concordance was poor. This may be due to the fact that amikacin is not affected by the action of most aminoglycoside modifying enzymes [35]. Previous studies predicting antimicrobial susceptibility from WGS data have reported sensitivities of 96% and 99% against phenotypic AST as a benchmark [20,21], compared with an overall sensitivity of 76% in this inter-laboratory study. It should be noted however that some of the data used in this study were purposefully low quality and some of the clinical isolates were deliberately chosen to be difficult to characterise as our aim was to identify the contributors to discordant results reported between participants working on the same data in order to provide useful recommendations.

We found three stages of analysis that contributed to discrepancies in predictions: The quality of the sequence data used, the bioinformatic methods (choice of database or software used) and the interpretation of those results. Where single gene calling is required (e.g presence of a carbapenemase) results are mainly affected by sequence quality. However, once multiple genes are involved, all three analytical issues become important. We found the largest contributors to discrepant results between the gene variants reported in each sample and the phenotypic resistance predictions were the sample read depth and the choice of reference resistance gene database. Samples must be sequenced to a sufficient depth as well as sufficient coverage for the expected size of the genome, usually inferred by mapping to a suitable reference genome, of at least above 90% to ensure even coverage. Based on our own experience and these results, we recommend 30X as a lower limit. This also tends to be a default setting for many read assembly tools but generally most samples should have a higher depth of coverage than this for meaningful prediction. Some participants did flag that they would not normally analyse the low coverage samples (<30X, samples B-1, E and G) and if those samples are excluded from this analysis sensitivity in comparison to phenotypic AST rises from 76% to 98%. As long as the sequence data produced is of sufficient depth and quality (e.g. current Illumina error rates) we have observed the choice of sequencer and DNA library preparation method has a small effect on closely related gene variants but little discernible effect on the inference of resistance phenotype.

Some participants ran the same set of read data against different reference databases and merged the results which led to different gene variants being reported at the same loci. We also found reference sequences in different databases for same gene variant can differ by 15% nucleotide identity (*bla*_IMP-1_ in CARD and ARG-ANNOT). If precise identification of gene variants is required, we would strongly recommend avoiding this as it effectively leads to ‘double-dipping’ using the same reads. Multiple reference databases could be used but after screening for reads that have already been assigned a hit against one of the databases. This would avoid multiple different genes reported at the same genomic loci. However, it would be better to merge the different reference databases and remove the redundant sequences before comparisons are made against the test data. Sequence identity, and to lesser extent coverage cut-offs, should be kept high when comparing test data to a reference database. Based on this study we would recommend using sequence identity cut-off of at least 90%, in combination with an up to date reference resistance gene database. Although lowering of these thresholds does identify more candidate genes within a sample it did not improve concordance with phenotypic AST results in this study.

There is an overwhelming need for a standardised, centralised database that integrates the current knowledge base for linking genotype with resistance phenotype and is not linked to a single research group, as previously suggested [10]. There is also a growing need regarding computational reproducibility [36,37]. This would deal with many of the issues we have raised, such as which sequences to include and what gene nomenclature to use. With strict version control, such a resource would allow greater integration of results and be an invaluable tool for larger epidemiological studies. Currently, databases are being built for organisms such as for *Mycobacterium tuberculosis*, though this is a less challenging organism for genotype-phenotype predictions due to it being highly clonal and lacking an accessory genome [38,39]. A recent publication of a new protein-based database also obtained high concordance (98.4%) between genotype and phenotype for four food-borne pathogens [40]. However, for other clinically relevant organisms there are limited resources.

Participants in this study included a mixture of individuals and teams involved in AMR prediction in a variety of settings. A potential criticism is that we did not restrict these settings to those routinely predicting AMR phenotype for clinical use, meaning that some participants were attempting analyses they did not usually perform. However, the fact that AMR phenotype prediction from WGS is not yet routine in most clinical laboratories was the very reason for undertaking this study. Clinical laboratories at the moment do not have the tools or knowledge to make good phenotypic resistance calls from genotypic data. This is evident from the fact that two participants in this study did not report any phenotypic resistance predictions as they felt they could find no valid method for doing so. At this point in time many research laboratories use these methods to track specific resistance genes or one specific resistance mechanism, rather than building tools for the broad detection of AMR in bacteria. We found in this study that there was particularly low concordance between participants reporting sensitive isolates compared with phenotypic AST. The problem with the inference of phenotype from genotype is that the information is either not known at all or is expert knowledge restricted to single laboratories working on specific bacteria. In addition to this, although the identification of the presence of genes is performed in a systematic way, the prediction of resistance is still performed in an *ad ho*c manner by scientists and therefore subject to user error given the same set of genes. Once again *M. tuberculosis* is providing the first example of the need for a defined decision tree when working from the presence of genes to the prediction of phenotypic drug resistance [41]. Interpretation and reporting of this genotypic data will need to be subjected to the same level of scrutiny as current tests if it is to form part of an accredited laboratory service within the healthcare service.

A limitation of this study is that we focused on the use of short read sequence data which produces sequences far shorter than the length of genes being identified. However, we feel this is more reflective of the WGS data that is more routinely generated in clinical laboratories at this point in time. If these short reads need to be assembled into longer contiguous sequences and we found the use of an actively developed short read assembler to be essential for this. Web-hosted tools that provide a “black box” solution to assembly and identifying resistance from uploaded WGS data should be avoided if possible, because of the lack of interpretability. Tools are needed which are open source, designed for purpose and can be subjected to thorough troubleshooting when erroneous results arise [42]. To this end, permanently employed bioinformaticians are required who can provide expert interpretation of the results and update approaches as necessary. In this study, tools that either require assembled contigs (ABRicate) and those that take unassembled short reads (SRST2 and ARIBA) were capable of producing very similar results with no notable effects alone on the predication of phenotypic resistance. This hold promise for rapid phenotypic predictions as genome assembly is one of the largest bottlenecks in computational analysis time.

Other limitations of this study include our focus on acquired genes rather than point mutations or many of the other resistance mechanisms found in bacteria (e.g. target site modifications and efflux pumps). We also only required reporting on categorical resistance predictions. Furthermore, because our focus was on WGS, although we validated AST at two independent laboratories we did not investigate potential variability and discordance in phenotypic prediction. More work needs to be done on the prediction of MICs from WGS data before it can be implemented in laboratories. This will be aided by more systematic reporting of accompanying MIC data when making WGS data available.

In conclusion, we have identified some of the current the key contributors to discrepancies in predicting AMR-associated genes and phenotypes from bacterial isolate WGS data. We have provided recommendations for improving the current reporting of results. Even after accounting for poor sequence data we found that the current public methods, in particular databases, are not adequate ‘off-the-shelf’ tools for the prediction of AMR from bacterial WGS data as a universal clinical test at this point in time.

## Supporting information

Excel spreadsheet template used by study participants to communicate results from each analysis.

Supplementary methods outlining individual study participant pipelines used in data analysis.

Contains Figure S1 which shows the presence of all AMR-associated genes in each sample by each participant.

## Abbreviations

AMR: Antimicrobial resistance
ARG-ANNOT: Antibiotic resistance gene-annotation
ARIBA: Antimicrobial resistance identification by assembly
AST: Antimicrobial susceptibility testing
CARD: Comprehensive Antibiotic Resistance Database
EUCAST: The European Committee on Antimicrobial Susceptibility Testing
GOSH: Great Ormond Street Hospital
NCBI: The National Center for Biotechnology Information
SRST2: Short read sequence typing 2
UHG: University Hospital Galway
WGS: Whole-genome sequencing

## Declarations

### Ethics approval and consent to participate

All investigations were performed in accordance with the hospitals’ research governance policies and procedures. No specific ethical approval was required, as no patient samples or identifiable data were used. The project was registered as a research study.

### Consent for publication

Not applicable.

### Availability of data and materials

The datasets generated and/or analysed during the current study are available in the European Nucleotide Archive at https://www.ebi.ac.uk/ena/data/view/PRJEB3451.

### Competing interests

ACP and AVB are employees of bioMérieux, a company developing, marketing and selling tests in the infectious disease domain. The company had no influence on the design and execution of the clinical study neither did the company influence the choice of the diagnostic tools used during the clinical study. The opinions expressed in the manuscript are the author’s which do not necessarily reflect company policies. All other authors declare that they have no competing interests and have performed the work in an individual capacity.

MJE and NW are members of PHE’s AMRHAI Reference Unit which has received financial support for conference attendance, lectures, research projects or contracted evaluations from numerous sources, including: Accelerate Diagnostics, Achaogen Inc., Allecra Therapeutics, Amplex, AstraZeneca UK Ltd, AusDiagnostics, Basilea Pharmaceutica, Becton Dickinson Diagnostics, bioMérieux, Bio-Rad Laboratories, BSAC, Cepheid, Check-Points B.V., Cubist Pharmaceuticals, Department of Health, Enigma Diagnostics, ECDC, Food Standards Agency, GlaxoSmithKline Services Ltd, Helperby Therapeutics, Henry Stewart Talks, IHMA Ltd, Innovate UK, Kalidex Pharmaceuticals, Melinta Therapeutics, Merck Sharpe & Dohme Corp., Meiji Seika Pharma Co., Ltd, Mobidiag, Momentum Biosciences Ltd, Neem Biotech, NIHR, Nordic Pharma Ltd, Norgine Pharmaceuticals, Rempex Pharmaceuticals Ltd, Roche, Rokitan Ltd, Smith & Nephew UK Ltd, Shionogi & Co. Ltd, Trius Therapeutics, VenatoRx Pharmaceuticals, Wockhardt Ltd and WHO

### Funding

This work was supported by the UK National Measurement System and the European Metrology Programme for Innovation and Research (EMPIR) joint research project [HLT07] “AntiMicroResist” which has received funding from the EMPIR programme co-financed by the Participating States and the European Union’s Horizon 2020 research and innovation programme. Andreu Coello Pelegrin received funding from the European Union’s Horizon 2020 research and innovation programme “New Diagnostics for Infectious Diseases” (ND4ID) under the Marie Sklodowska-Curie grant agreement N° 675412. These funding bodies had no influence on the design of the study and collection, analysis, and interpretation of data and in writing the manuscript.

### Authors’ contributions

RMD, DMO’S, KAH and JFH conceived and designed the study. SDA, SB, TC, ACP, MC, EDB, MJE, EM, YM, TPTN, JP, LPS, RAS, AvB, LvD and NW all performed the initial participant analyses, and are listed in alphabetical order. Only those on the author list from participating institutions contributed to this analysis. RMD performed all secondary analyses and drafted the manuscript with assistance from DMO’S, JMG, JFH and KAH. All authors read and approved the final manuscript.

## Acknowledgements

The authors thank the Biomedical Scientist teams for sample collection and processing. We also thank the Pathogen Informatics Group at the Wellcome Sanger Institute for their contributions to the study.

## Supplementary material

**Additional File 1:** Excel spreadsheet template used by study participants to communicate results from each analysis.

**Additional File 2:** Supplementary methods outlining individual study participant pipelines used in data analysis.

**Additional file 3:** Contains **Figure S1** which shows the presence of all AMR-associated genes in each sample by each participant. Genes are organised and coloured by the class of antibiotics they are associated with resistance, or if they are associated with the efflux of multiple classes of antibiotics.

Participants

## Notes

#### Summary of Updates

Additional text and declarations added to clarify that researchers participated in this study in an individual capacity. Only those named in the author list assisted in any analysis from their affiliated institution. Discussion was revised to reflect further on possible study limitations and two recent studies were also additionally referenced (41 and 42).

https://www.ebi.ac.uk/ena/data/view/PRJEB34513

