## Supplementary methods outlining individual study participant pipelines used in data analysis. for "Discordant bioinformatic predictions of antimicrobial resistance from whole-genome sequencing data of bacterial isolates: An inter-laboratory study"

Each laboratory was asked to provide a description of the pipeline used within this study. The submitted methods sections are included below:

### Laboratory 1

Raw reads were checked for quality using FastQC, trimmed/assembled/corrected/reassembled using shovill pipeline (see https://github.com/tseemann/shovill). Species ID was achieved with the raw reads using Kraken-HLL (see https://github.com/fbreitwieser/krakenhll) and Bracken (for abundances, see https://github.com/jenniferlu717/Bracken). AMR analysis involved searching the assembled contigs CARD database with RGI tool and ResFinder/ARG-ANNOT databases with the software tool c-SSTAR (see https://github.com/chrisgulvik/c-SSTAR ).

### Laboratory 2

Raw reads trimmed using Trimmomatic and Trim Galore. Reads assembled using SPAdes. Resistance genes detected using abricate with the CARD database on assemblies. BLASTs on 16S and other genes and mlst-master used to identify species.

### Laboratory 3

#### Read QC, Assembly

10 paired end fastq files were downloaded from the GOSH server. Reads were initially screened for quality using FastQC (https://www.bioinformatics.babraham.ac.uk/projects/fastqc/)

Raw reads, were assembled using UniCycler (Wick *et al.*, 2017) in "Illumina-only" assembly mode. UniCycler uses SPAdes' built-in read correction module. Here we used SPAdes version 3.10.1 (Bankevich *et al.*, 2012) and generate a SPADes assembly graph. UniCycler then performs additional assembly improvement steps through identifying multiplicity of contigs, scaffolding, overlap removal and bridging (see Wick *et al*. for details). The resultant assemblies were visualized in Bandage (Wick *et al.*, 2015) to check overall quality and merge overlapping graphs.

#### Species Assignment

Taxonomic assignment was performed using two approaches. The first utilized the raw sequencing reads, without any assembly step. Raw reads were screened using the distance based tool

MASH(Ondov *et al.*, 2016), which implements the minHASH k-mer matching algorithm. MASH distances were calculated for each of the 10 genomes against an archive of RefSeq genomes (Pruitt *et al.*, 2012), release 70, sketched using $k$=21 and $s$=1000. The closet matching reference was selected based on the proportion of matching k-mers.

In addition, and to validate the assignments using MASH, the *de novo* assemblies were uploaded to WGSA (wgsa.net), which provides a rapid taxonomic assignment.

**AMR Gene Detection**

Denovo assemblies were uploaded to 2 independent reference databases for AMR gene identification: CARD (Jia *et al.*, 2017) and ResFinder (Zankari *et al.*, 2012), the latter was run with a %ID threshold of 90% and a selected minimum length of 80%. Resulting output files were downloaded and inspected.

**Assigning Resistance Phenotype**

Resistance phenotypes were considered in the context of the presence, absence and co-occurence of different AMR genes. We took a conservative approach corresponding to the 'strict' classification in CARD, calling a resistance gene present if there was >80% identity to the reference. We operated on a precautionary principle (appropriate for clinical work, where WGS could be used to triage samples) and called a gene as present even in cases where not all of the reference gene was covered (e.g. AMRIL_2, with 22% coverage of SHV-156).

The CARD reference database ontology terms were used to relate resistance genes to resistance to specific antibiotics. In addition to using this ontology, we discussed the following heuristic reasoning for our phenotype classifications based on our existing knowledge:

**Ciprofloxacin** (fluoroquinolone): a combination of specific genes (e.g. *patA* and *patB, Qnr1*) and mutations (e.g. in *gyrA* and *gyrB*).

**Gentamicin** and **amikacin** (aminoglycoside): aminoglycoside resistance is conferred by specific genes (e.g. *AAC* family). AAC(3)-II confers resistance to gentamicin, but AAC(6')-I is additionally required for resistance to amikacin.

**Cefotaxime** (beta-lactam): many different beta-lactamases, with resistance correspondingly growing additively with the number of genes present.

### Laboratory 4

Ariba (https://github.com/sanger-pathogens/ariba) was used to identify antibiotic resistance genes and MLST sequence types by running local assemblies.

Since ariba does not allow species identification, kraken (https://ccb.jhu.edu/software/kraken/; https://genomebiology.biomedcentral.com/articles/10.1186/gb-2014-15-3-r46) was used to identify the species before ariba was run to identify the sequence type. The kraken analysis is based on a database from the following publication: Browne at al. Nature 2016, https://www.nature.com/articles/nature17645

To identify antibiotic resistance genes, the two following databases were used: The Comprehensive Antibiotic Resistance Database "CARD" version 2.0.1 (https://card.mcmaster.ca/home; https://www.ncbi.nlm.nih.gov/pubmed/23650175) and Antibiotic Resistance Gene-ANNOTation

“ARG-ANNOT” (http://en.mediterranee-infection.com/article.php?laref=283&titre=; https://www.ncbi.nlm.nih.gov/pubmed/24145532). Databases were downloaded using ariba’s getref command on June 27th 2018. Ariba was run using standard settings and results were checked manually. Disparate results between the two databases are specified in the comment column, only the hit with the best identity was kept. Possible contaminations were highlighted in the commend section when the sequence was incomplete and its read coverage deviated from the coverage of other contigs (column ctg_cov in the original ariba reports). Only hits of genes conferring resistance to aminoglycosides, beta-lactams and fluoroquinolones were listed in the Excel file “AMRIL_WGS_reporting”. All hits were kept in the original ariba report files (“AMRIL_*_CARD_report.tsv”; “AMRIL_*_argannot_report.tsv”)

Ariba does not predict AMR phenotypes. Therefore, these fields have been left empty.

### Laboratory 5

Fastq files quality was checked with FastQC. Then, reads were assembled using the A5-miseq pipeline, and resulting contigs were analysed with the Resistence Gene Identifier tool, using as reference the Comprehensive Antimicrobial Resistance Database. In detail: FastQ files were downloaded and analyzed with the FastQC program (v0.11.7) by Babraham bioinformatics. Next, reads are assembled using the A5_miSeq pipeline (v20160825), using standard parameters. A short bash script is used to accelerate the process. ID was obtained uploading the assembled contigs to the online KmerFinder tool (v2.5) from CGE. Conda is used for the posterior analysis, using the RGI (v 4.0.3) software. Also, last version of the CARD database (v 2.0.2) is used. Another bash script is used to run RGI. Results with a Cut-off marked as "Loose" were not took into account.

### Laboratory 6

We used BioNumerics version 7.6.3 with E.coli plug in tools from CGE (ResFinder). K-mer finder was used to determine bacterial species http://www.genomicepidemiology.org/. We did not search for *gyrA* and *parC* mutations (quinolone resistance) in these sequences.

### Laboratory 7

Reads were trimmed using trimmomatic. Species classification was performed using centrifuge. Resistance genes were found using SRST2. Resistance to specific drugs were predicted by assessing the literature

**Read trimming**

Program: trimmomatic
Version: v0.38
Program call:
trimmomatic PE -threads 1 -phred33 AMRIL_1_R1_001.fastq.gz AMRIL_1_R2_001.fastq.gz -baseout AMRIL_1 LEADING:3 TRAILING:3 SLIDINGWINDOW:4:20 MINLEN:36

**Species classification**

Program: Centrifiuge
Version: 1.0.3-beta
Database: ftp://ftp.ccb.jhu.edu/pub/infphilo/centrifuge/data/p+h+v.tar.gz

Program call:
centrifuge -x p+h+v -1 AMRIL_1_1P -2 AMRIL_1_2P > AMRIL_1.log
centrifuge-kreport x p+h+v AMRIL_1.log > AMRIL_1.kreport

**Drug resistance gene detection**
Program: SRST2
Version: 0.2.0
Database: https://github.com/katholt/srst2/blob/master/data/ARGannot_r2.fasta
Program call:
srst2 --input_pe AMRIL_1_R*.fastq.gz --forward _R1_001 --reverse _R2_001 --output test --gene_db ARGannot_r2.fasta

### Laboratory 8

**Table 1|** List and description of tools utilized in this study.

| Name | Associated Workflow (Fig. 1) | Description |
|---|---|---|
| Conda | N/A | Package, dependency and environment management for multiple programming languages. (https://conda.io/docs/) |
| Snakemake [1] | N/A | Python based workflow management system. (https://snakemake.readthedocs.io/en/stable/) |
| Sickle | trim | C based program for adaptive trimming of FASTQ files [according to quality]. (https://github.com/ucdavis-bioinformatics/sickle) |
| Unicycler [2] | assembly | Assembly pipeline for bacterial genomes from NGS reads; acts as an optimizer for SPAdes. (https://github.com/rrwick/Unicycler) |
| SPAdes [3] | assembly | Python based genome assembler with built-in read error correction. (http://cab.spbu.ru/software/spades/) |
| ABRicate | amr | Perl based program for screening of assembled genomes [contigs] for antimicrobial resistance or virulence genes [database specific]. (https://github.com/tseemann/abricate) |
| Kraken [4] | taxonomy; taxonomy-report | C++, Perl and Shell based program for taxonomic classification from sequences files [individual reads, assembled genomes etc]. (https://github.com/DerrickWood/kraken) |

#### Workflow Overview

1. FASTQ reads are trimmed using **sickle** yielding trimmed paired reads 1 and 2, and another trimmed reads file containing the single, unpaired reads.

```
sickle pe -f read_1 -r read_2 -t sanger \
                    -o trimmed_read_1 -p trimmed_read_2 -s trimmed_read_singles
```

2. Trimmed FASTQ (1, 2 and singles) are then assembled using the Unicycler optimizer for SPAdes. The Unicycler optimizer produces a single ‘assembly.fasta’ file (in place of the ‘contigs.fasta’ and ‘scaffolds.fasta’ default output from SPAdes).

```
unicycler-runner.py -1 trimmed_read_1 -2 trimmed_read_2 -s trimmed_read_singles \
                    -t <n_cpus> --mode normal -o output_folder
```

3. Antimicrobial resistance (AMR) is then predicted using the ABRicate tool, along with the ResFinder database (note that resfinder is selected over other AMR databases as it only focuses on acquired AMR genes, not chromosomal point mutations). Report is generated as ‘.csv’ output.

```
abricate --db=resfinder assembly.fasta --csv > amr.csv
```

   The output of ‘abricate’ is a list of predicted acquired AMR genes. These genes correspond to specific drug-class resistance (i.e. aminoglycosides) as indicated in the ResFinder database. These gene-class associations are used to predict the AMR phenotype of samples to specific drugs (by identifying the class associated with each drug, i.e. ciprofloxacin → fluoroquinoline).

4. Taxonomic predictions are performed by Kraken (as well as report generation). Kraken utilizes a local database, downloaded from the Kraken repository (the standard database). Taxonomic prediction file is generated by:

```
kraken --preload --db <DATABASE_FOLDER> --fasta-input assembly.fasta --threads
<n_cpus> > kraken.out
```

   And the subsequent [human-readable] report generated by:

```
kraken-report --db <DATABASE_FOLDER> kraken.out > kraken.txt
```

   The species selection is done by identifying the species with the largest percentage abundance in the sample.

5. The ‘all’ rule is a Snakemake specific workflow rule that indicates all the above rules (as indicated in Fig. 1) are required to be carried out.

### Laboratory 9

In-house program “qa_and_trim” calls Trimmomatic; nucleotides with a Phred score less than Q30 at the ends of the reads were removed with Trimmomatic. In house “kmerid” was used for kmer-based species identification (https://github.com/phe-bioinformatics/kmerid). Resistance gene detection was done using ‘Genefinder’, an in-house algorithm that uses bowtie2 to map sequence reads to reference sequences of interest and Samtools vs 0.1.18 to generate an mpileup file, which is then parsed for the rapid detection of sought sequences. Genes were called as present within a genome when detected with 100% coverage and >90% nucleotide identity to the reference gene.
