## Supplementary figures and images for "Discordant bioinformatic predictions of antimicrobial resistance from whole-genome sequencing data of bacterial isolates: An inter-laboratory study"

### Contains Figure S1 which shows the presence of all AMR-associated genes in each sample by each participant.

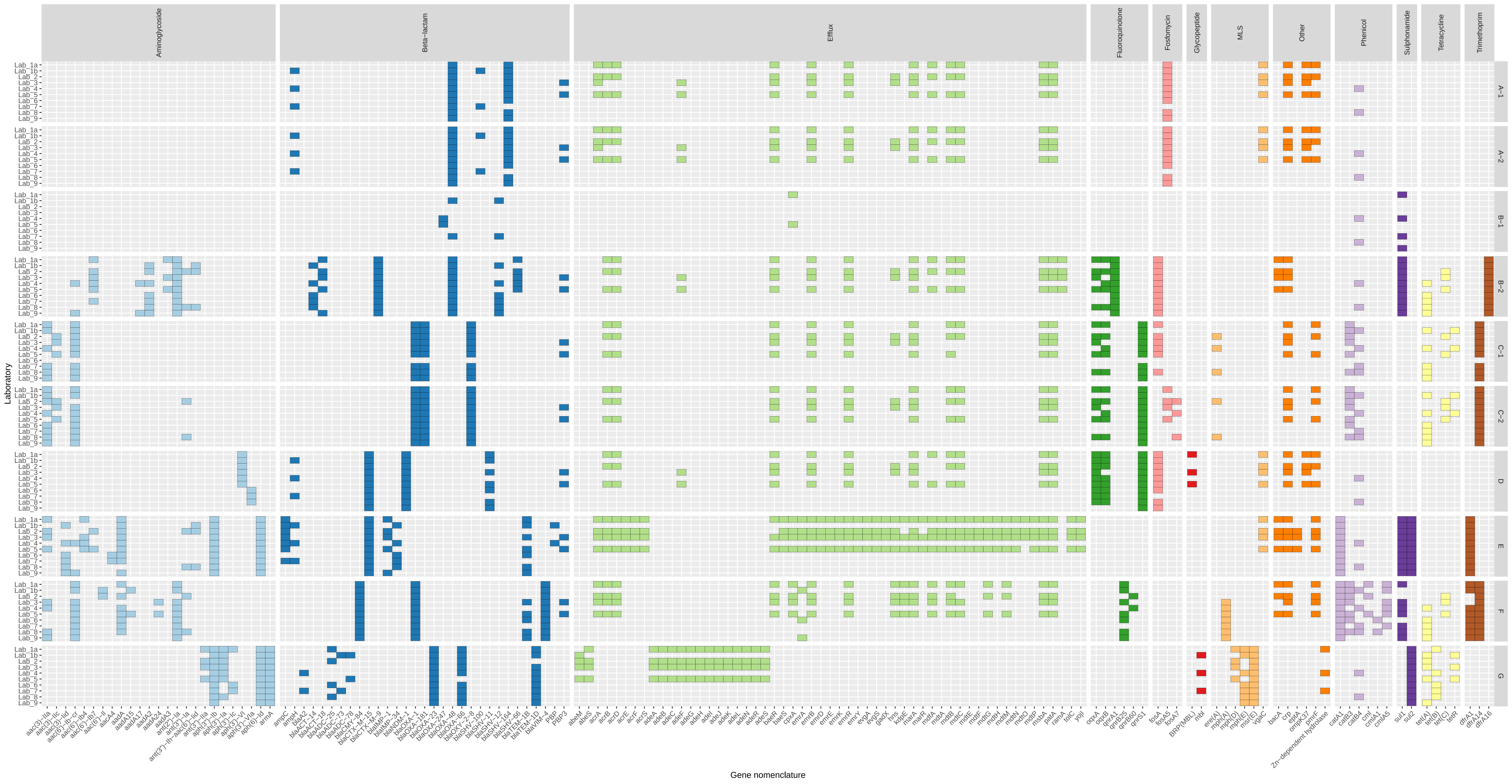
